# predatoR: an R package for network-based mutation impact prediction

**DOI:** 10.1101/2022.11.29.518310

**Authors:** Berk Gurdamar, Osman Ugur Sezerman

## Abstract

**Motivation:** Classification of a mutation is important for variant prioritization and diagnostics. However, it is still a challenging task that many mutations are classified as variant of unknown significance. Therefore, in silico tools are required for classifying variants with unknown significance. Over the past decades, several computational methods have been developed but they usually have limited accuracy and high false-positive rates. To address these needs, we developed a new machine learning-based method for calculating the impact of a mutation by converting protein structures to networks and using network properties of the mutated site.

**Results:** Here, we propose a novel machine learning-based method, predatoR, for mutation impact prediction. The model was trained using both VariBench and ClinVar datasets and benchmarked against currently available methods using the Missense3D datasets. predatoR outperformed 32 different mutation impact prediction methods with an AUROC value of 0.941.

**Availability:** predatoR tool is available as an open-source R package at GitHub (https://github.com/berkgurdamar/predatoR).

## 1 Introduction

Single nucleotide variations in the coding regions of the genome can alter the amino acid sequence of a protein and may results in change in the protein structure and function. Such variations are called missense variants, or single amino acid variants (SAVs), which are the most studied form of variations (Yates *et al*., 2014). SAVs may affect the protein stability and cause alteration in the active site, protein-protein interaction sites and thus affect the function of the protein (Tan *et al*., 2021).

Developments in the next-generation sequencing technologies have enabled identification of several pathogenic mutations and establishment of disease-causing mutation databases such as ClinVar (Landrum *et al*., 2014), OMIM (Hamosh *et al*., 2005) and HGMD (Stenson *et al*., 2014). These databases facilitated prompt diagnosis of patients if their variants are present in the database. However, majority of the variations are still classified as variant of unknown significance (VUS) due to low frequency of such variants and lack of information about the association of the variant with a certain phenotype. Especially in rare diseases, most of the variants in a patient are not present in the databases. Therefore, there is an imminent need for more accurate impact prediction methods, which will provide a cost-efficient and time-efficient solution in contrast to experimental methods (Chen *et al*., 2020).

In the last decade, several computational methods and in-silico tools have been developed for mutation impact prediction. Most mutation impact prediction tools exploit the sequence, structure, and thermodynamic information and use several machine learning-based methods for prediction. However, accuracies of the methods remain limited with high false-positive rates (Ancien *et al*., 2018).

Different approaches have been made, such as GERP++ (Davydov *et al*., 2010) and SIFT (Ng and Henikoff, 2003) which make predictions using multiple sequence alignment-based conservation measuring. On the other hand, methods such as CADD (Rentzsch *et al*., 2019) uses a machine learning model with multiple genomic features. Moreover, Rhapsody (Ponzoni *et al*., 2020) uses structural information along with sequence and thermodynamic-based information. Another approach is the metapredictors whose main approach is aggregating predictions of other predictors. BayesDel (Feng, 2017), REVEL (Ioannidis *et al*., 2016), and FATHMM-XF (Rogers *et al*., 2018) are such examples of metapredictors. Thus far, network formalization have not been used for mutation impact prediction. ProSNEx (Aydinkal *et al*., 2019) is the only method uses network formalization from protein structures, but the tool provides structure exploration not impact prediction.

Considering the downfalls of the aforementioned in-silico methods, we developed predatoR, a new machine learning-based method for mutation impact prediction. predatoR was implemented in R (R Core Team, 2021), and uses a network formalization approach that accepts protein structures as networks and uses 24 unique features for calculating the impact of a mutation. By putting all the factors that influence the impact of a mutation, we demonstrate that predatoR outperforms 32 different impact prediction methods currently in widespread use.

## 2 Materials and Methods

### 2.1 Overview

Main aim of this study was to build a new machine learning-based method for mutation impact prediction by converting protein structures into networks. For calculating network properties, PDB structures were converted into networks by calculating distances between atoms and assigning edges according to interatomic interaction distance cutoffs. Twenty-four unique features were taken into consideration. Machine learning models were trained using VariBench (Sasidharan Nair and Vihinen, 2013) and ClinVar (Landrum *et al*., 2014) datasets. Seven different machine learning algorithms were employed and the performance of the models were tested using the Missense3D (Ittisoponpisan *et al*., 2019) dataset. Performance of the novel method was compared with currently available 32 different mutation impact prediction methods by using the predictions on the Missense3D dataset. Details of this study are described in the following sections.

### 2.2 Data Collection

Neutral datasets (DS2, DS4, DS6, DS8, DS10, DS12, DS14, DS16, MMR and dbSNP), pathogenic datasets (DS3 and DS5) and training dataset of Rhapsody were downloaded from VariBench web server. Duplicated and conflicting mutations (the same mutation labeled as both pathogenic and neutral) were removed. The combined VariBench dataset contained 6983 neutral and 23563 pathogenic mutations. For increasing the number of neutral mutations and preventing class imbalance, benign and likely benign variants from ClinVar were also collected. Variants in the ClinVar datasets were only mapped on genomic positions. For converting genomic positions to PDB-based positions, VarMap (Stephenson *et al*., 2019) was used. From the output, PDB position containing variants were filtered and combined with the VariBench dataset. Duplicated, conflicting and synonymous mutations were removed, and the final dataset contained 48005 mutations (24496 neutral and 23509 pathogenic variants).

Experimental datasets of Missense3D were selected for validation of machine learning models and comparison with available impact prediction methods. 2 datasets were downloaded from Missense3D web server. The combined dataset contains 10229 different mutations (4652 neutral, 5577 pathogenic variants). Only UniProt-based positions of the mutations were available in the dataset. For converting UniProt positions to PDB-based positions, the PDBSWS (Martin, 2005) dataset was used. Variants with PDB-based positions were filtered and common variants between Missense3D dataset and merged ClinVar-VariBench dataset were removed. Final size of the Missense3D dataset decreased to 6554 mutations (3569 neutral, 2985 pathogenic variants).

### 2.3 Processing

For model training, we used 24 different features divided into 3 categories: Network-based, gene-based and amino acid-based. Network-based features were calculated by turning protein structures into networks from PDB structures. For this approach, PDB structures were downloaded by using the Bio3D (Grant *et al*., 2006) R package. Two different network building approaches were tested: Building network using only carbon alpha (Cα) atoms and all atoms in the structure. For each approach, networks were built using 2 different interatomic distance cutoffs: ≤ 7 angstrom (Å) and ≤ 5Å. According to the approach, edge lists were created by calculating distance between each atom. From the edge list, networks were created for each chain separately via igraph (Csardi and Nepusz) R package. These networks were used for calculating Eigen Centrality, Betweenness Centrality, Degree Centrality, PageRank Centrality and Shortest Path Centrality measures by using the igraph R package and node-specific Clique Score by using in-house scripting. For Shortest Path Centrality, sum of all shortest path lengths of a node were calculated and the sum of the shortest path lengths were used as a score of a node. For node-specific Clique Score, the number of edges between neighbors of a node was calculated for each node separately. For normalizing the scores, scores of network properties were turned into Z-scores for each chain separately for preventing bias between high and low number of atom containing structures. Z-scores of the Cα atoms were accepted as the scores of the positions.

For gene-based features, each mutation was annotated according to its corresponding gene identifier. However, not all the mutations in the datasets contained gene identifier information. For completing the missing gene identifiers, Ensembl BioMart (Smedley *et al*., 2009) web server was used for mapping PDB-chain IDs to associated gene identifiers. Nine different gene-based features were used. “pLoF Metrics by Gene” dataset was downloaded from gnomAD (Karczewski *et al*., 2020) web server. From this dataset Missense Z-scores, Synonymous Z-scores and pLI scores were filtered. All human KEGG pathways and associated genes were downloaded by using EnrichmentBrowser (Geistlinger *et al*., 2016) R package and number of pathways associated with each gene were calculated. Ensembl BioMart web server was used for collecting biological process domain related and experimental evidence-coded (EXP, IDA, IPI, IMP, IGI, IEP, HAD, HMP and HEP) GO terms and gene associated GO term numbers were calculated. Disease-Gene association dataset was downloaded from DisGeNET (Piñero *et al*., 2017) database and gene related disease numbers were computed. Moreover, Genic Intolerance (Petrovski *et al*., 2013) scores and gene essentiality scores from OGEE v3 (Gurumayum *et al*., 2021) were downloaded. The median TPM expression dataset of the GTEx portal (Lonsdale *et al*., 2013) was collected and median gene expression values across 54 different tissue types were calculated. Each feature was annotated/calculated according to the associated gene name in the dataset. If there were multiple genes associated with the same PDB-chain ID, the gene having higher gnomAD scores across all 3 metrics was selected.

Nine different amino acid-based features were used. Reference amino acids and the mutant amino acids were used as 2 categorical features. Accessible Surface Area (ASA) and Hydrophobicity Scale values were collected from aaSEA (D.M, 2019) R package. ASA and Hydrophobicity Scale of reference amino acids, mutant amino acids and the difference between mutant and the reference amino acids were calculated. Lastly, BLOSUM62 scores between reference and the mutant amino acids were also used as another feature.

### 2.4 Model training/testing and comparison

After the processing step (Fig. 1A), the size of the dataset decreased to 34968 mutations (17469 neutral, 17499 pathogenic variants) due to missing information. The dataset was split into 90% training and 10% testing datasets with conserving the proportions of pathogenic and neutral variants. Impact prediction models were built using 7 different machine learning algorithms, XGBoost, Naïve Bayes, Random Forest, Elastic Net, GLM, GBM and Adaboost (Fig. 1B). Parameters were optimized through 3 repeated 10-fold cross-validation and for each model, same training-testing datasets were used. For method comparison, the Missense3D dataset went through the same processing steps, as described above (Fig. 1C). The final size of the dataset was 4403 mutations (1803 neutral, 2600 pathogenic variants) after the processing step. For annotating the Missense3D dataset with available impact prediction methods, ANNOVAR (Wang *et al*., 2010) was used. However, ANNOVAR uses genomic positions as an input. For converting PDB-based positions to genomic positions, TransVar (Zhou *et al*., 2015) was used. Impact prediction scores of 31 different methods were annotated onto the dataset via ANNOVAR using the DBNSFP 4.2a database. We also collected the pathogenicity predictions from Rhapsody on the Missense3D dataset by using the Rhapsody web server.

**Fig. 1.**
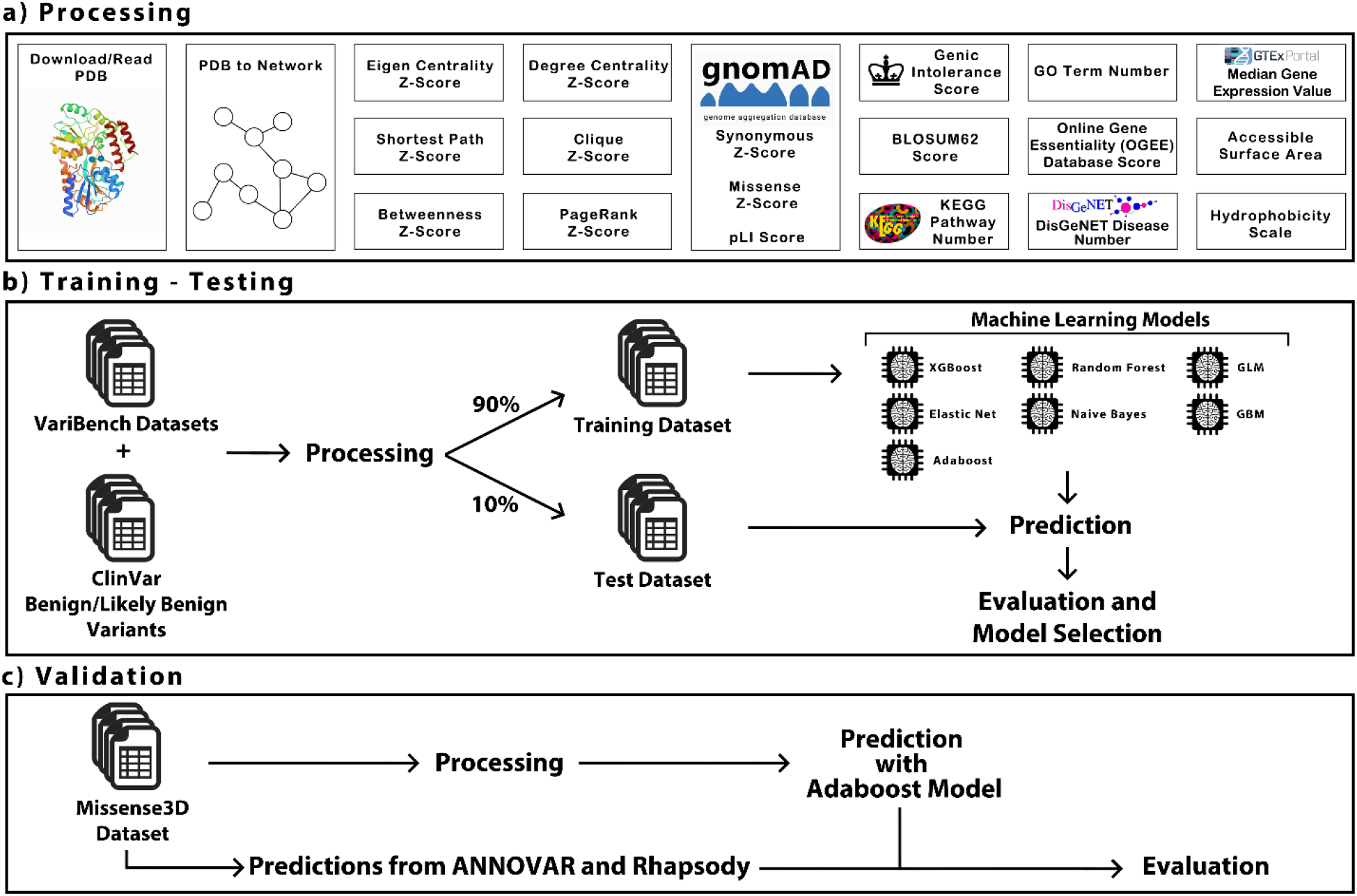
Overall workflow of the study. **(A)** Simplified processing steps of predatoR that were done on the training, testing and validation datasets. **(B)** Workflow of machine learning model training, testing and evaluation. **(C)** Simplified workflow of comparison between predatoR and available 32 different impact prediction methods with using Missense3D dataset.

## 3 Results

### 3.1 Performances of impact prediction models

Performance of the machine learning models were evaluated using ROC curves and AUROC curve values via testing dataset (Fig. 2). According to the AUROC curve values, for all atoms and Cα atoms approaches, 7Å-Adaboost model (AUROC: 0.9762) and 5Å-Adaboost model (AU-ROC: 0.9808) showed the best performance respectively. These two models were selected as final models and used for comparison with available impact prediction methods.

**Fig. 2.**
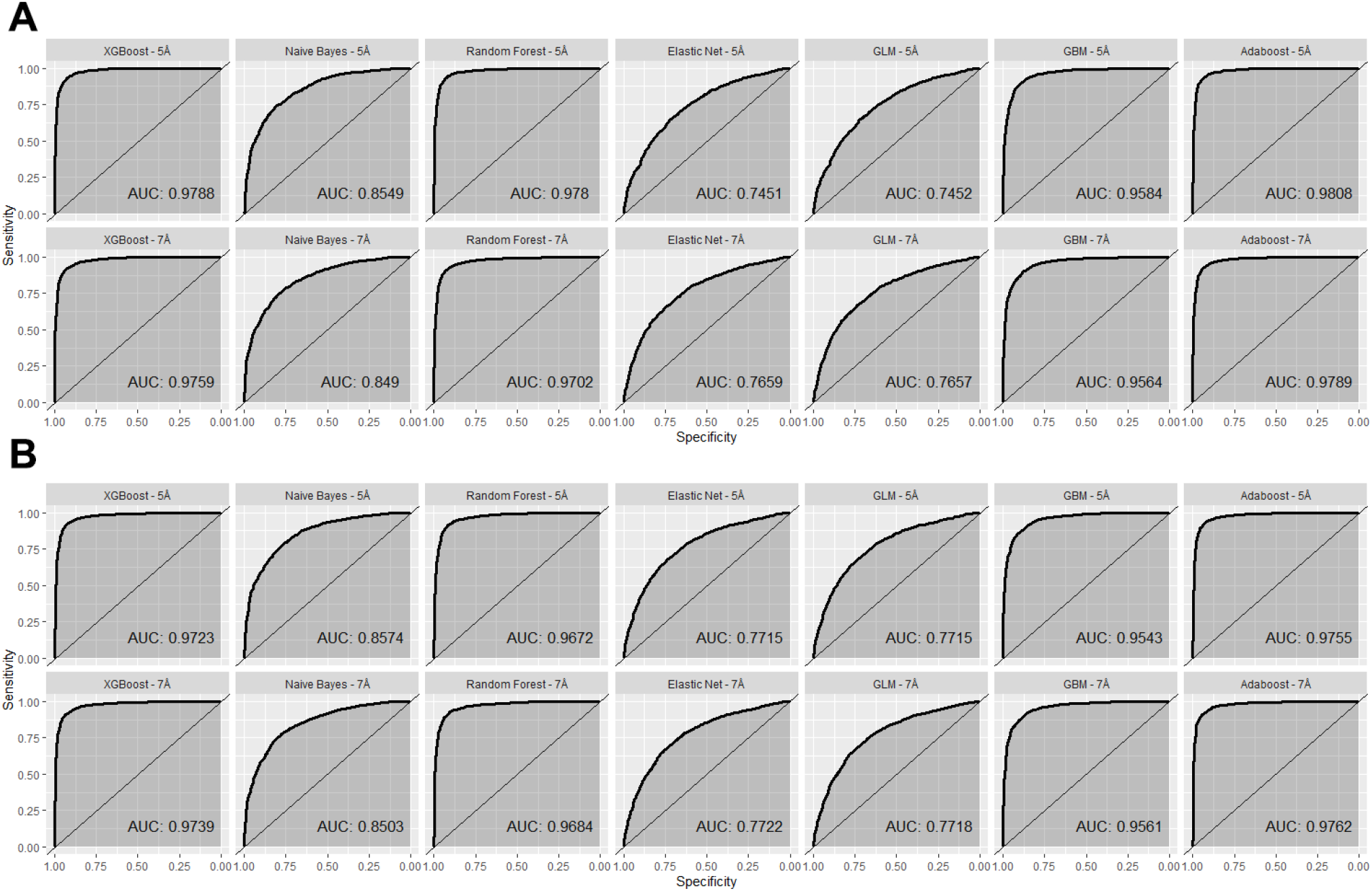
Performance of the machine learning models on the test dataset with Cα atoms approach **(A)** and all atoms approach **(B)**.

### 3.2 Comparison with other methods

Scores of 31 different impact prediction methods were annotated for the Missense3D dataset via ANNOVAR and predictions of Rhapsody were collected using their web server. The performance of all the prediction methods were visualized and evaluated using ROC curves and AUROC curve values (Fig. 3). According to the AUROC curve values, 7Å-all atoms approach used Adaboost model outperformed all other prediction tools with an AUROC curve value of 0.941. Feature importances of the final models represented in the Fig. 4.

**Fig. 3.**
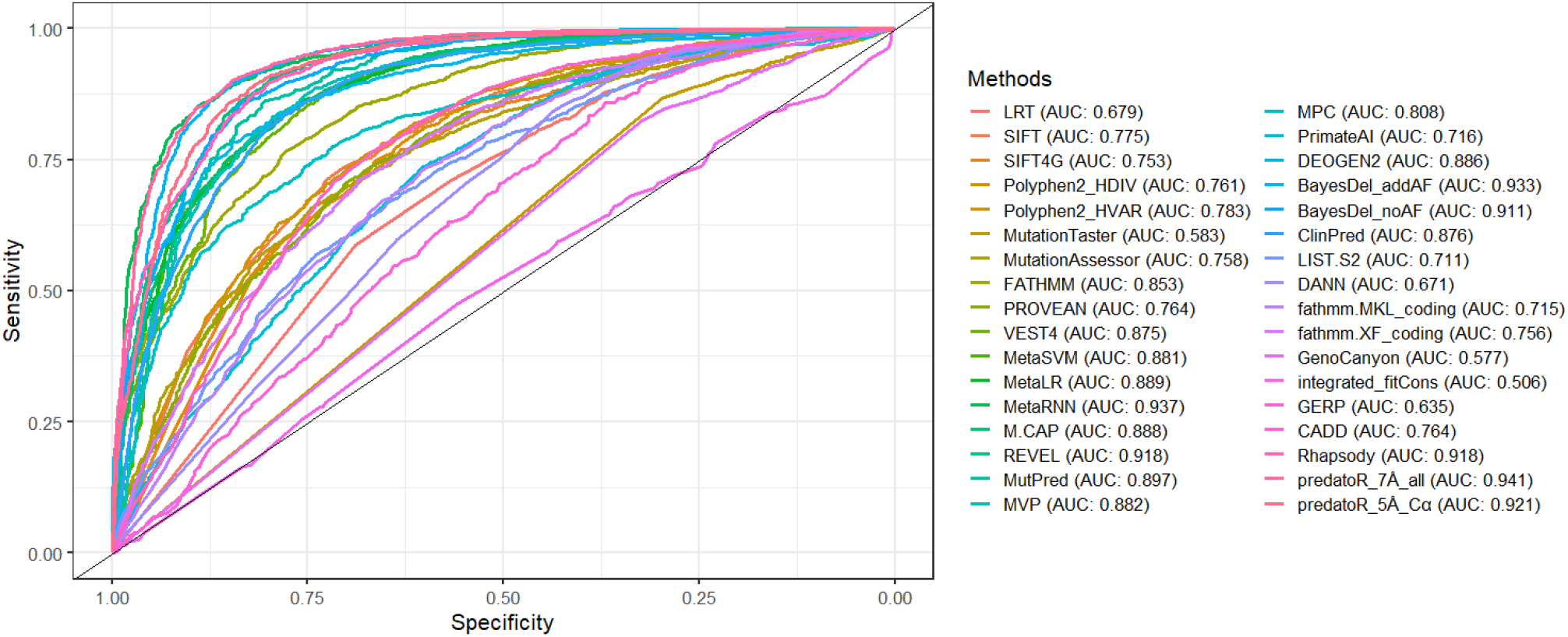
ROC curves and AUROC curve values of predatoR and 32 available impact prediction methods. ROC curves are based on the predictions on Missense3D dataset.

**Fig. 4.**
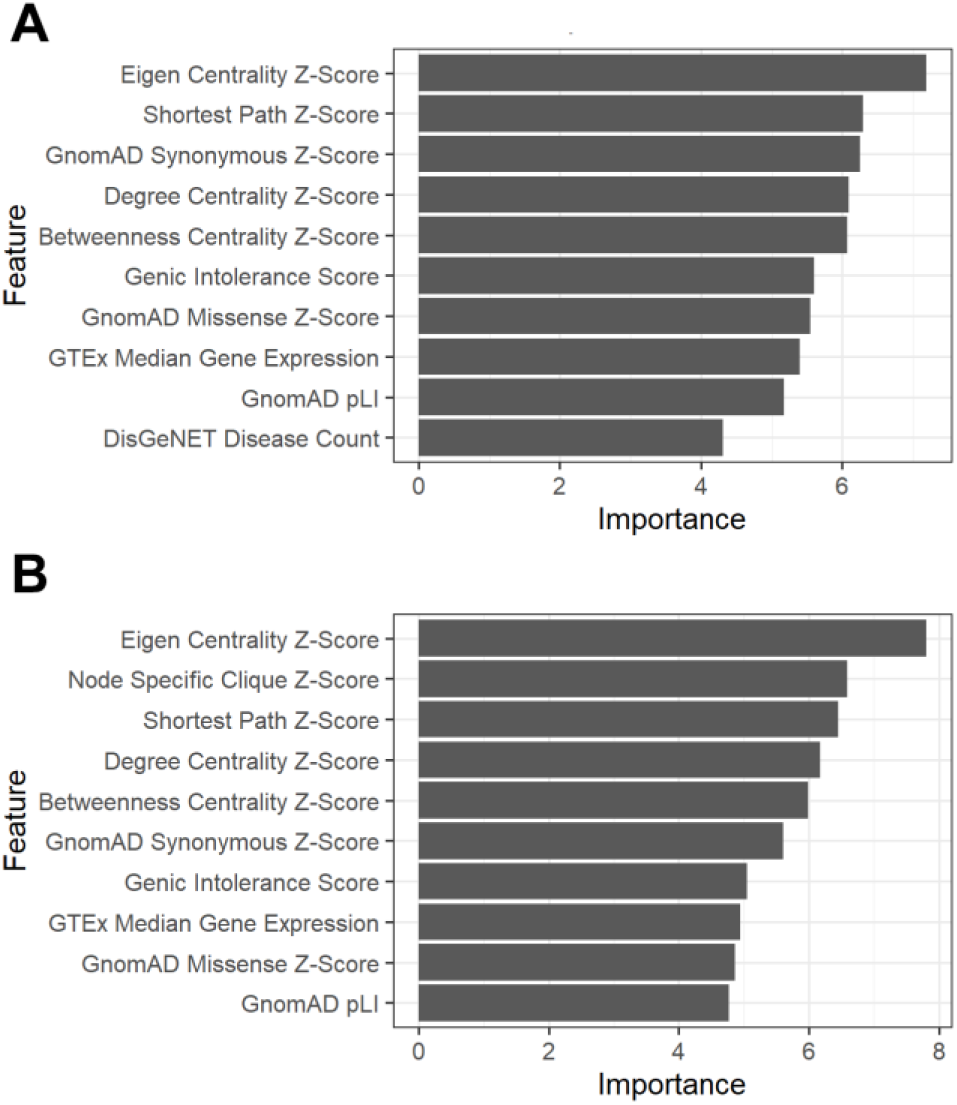
Feature importance plots of the 10 most important feature of the final models. (A) 7Å-all atoms approach used Adaboost model. (B) 5Å-Cα atoms approach used Adaboost model.

### 3.3 The predatoR R package

The predatoR package takes an input with 5 mandatory and 1 optional argument, PDB ID, chain ID, position (PDB-based), reference and mutant amino acids with 3 letter codes, and gene identifier (optional). If a gene identifier is not provided in the input, associated gene identifier is found using as described before. Package contains 2 models for prediction, 7Å-all atoms approach used Adaboost model and 5Å-Cα atoms approach used Adaboost model. User can decide which model they are going to use. Each utility function of the package can be used separately if necessary. Also, predatoR can be used for exploratory purposes, such as networks can be built with using different cutoffs. In exploratory purposes, all 24 features annotated to the input dataset and returned as the output. However, predictions cannot be made using a cutoff other than 5Å and 7Å. The predatoR package can be installed from GitHub (https://github.com/berkgurdamar/predatoR).

## 4 Discussion

In this study, we propose a new machine learning-based method for analyzing protein structures as networks and making predictions on the impact of a mutation with using 24 different unique features. Creating a network model from a protein structure is a novel and promising approach for mutation impact prediction. One of the advantages of this approach is that it uses structural information as network properties and takes less time, needs less computational power than thermodynamic simulations. Two different network formalization approaches (all atoms and Cα atoms) were tested for network building. For setting edges between atoms, two different interatomic distance cutoffs (5Å and 7Å) were tested. According to the results, models built with 7Å cutoff showed better AUROC curve values in most of the models on test dataset. Moreover, final model that uses all atoms approach showed better performance than the Cα atoms on the Missense3D dataset. This shows that, considering more interactions from the protein structure have positive effect on the prediction performance. Despite the better performance of all atom approach, Cα approach can build the networks and make predictions faster due to the fewer number of atoms used in the networks.

Two different models from two different network formalization approaches were selected as final models and compared with 32 different mutation impact prediction methods with using Missense3D datasets. 7Å-all atom approach used model outperformed all 32 other methods with an AUROC curve value of 0.941. Even though 5Å-Cα atom approach used model could not outperformed all the other methods, it is still performed well and showed better AUROC curve value compared with most of the methods. Therefore, it is included in the predatoR package and user can decide which model to use for prediction. In addition, exploratory analysis can be done by using predatoR. Main aim of exploratory analysis is that new models and new approaches can be made with using different interatomic interaction cutoffs or with combination of different features. We hope the community benefits from exploratory analysis.

For model training and testing, VariBench datasets were selected. However, after combining VariBench datasets, we found that most of the datasets contain pathogenic variants and number of neutral variants containing datasets are few or the datasets contain few number of variations. In addition, different datasets have conflicting labels (pathogenic and neutral) for the same mutations and those mutations had to be removed and it resulted in a decrease in the number of mutations. For preventing class imbalance and increase the number of neutral variants, benign and likely benign variants from ClinVar were used. Still, many mutations in the ClinVar dataset were filtered after the conversion from genomic coordinates to PDB positions due to the missing PDB annotations. With an update in the available databases and increase in the number of datasets, better models and methods can be built for mutation impact prediction and more advancements can be done in our field.

In summary, we developed a new machine learning-based method, predatoR, for mutation impact prediction by converting protein structures into networks and making prediction with using 24 different unique features. Apart from performing impact prediction, exploratory analysis also can be done. Networks can be built, and network properties can be calculated with desired thresholds and further analysis can be done. predatoR is presented as an R package and designed as user-friendly as possible.

## Funding

European Joint Programme on Rare Diseases (EJP RD) provides B.G. with a monthly scholarship.

### Conflict of Interest

none declared.

## Notes

### Competing Interest Statement

The authors have declared no competing interest.

